# PGxCorpus: a Manually Annotated Corpus for Pharmacogenomics

**DOI:** 10.1101/534388

**Authors:** Joël Legrand, Romain Gogdemir, Cédric Bousquet, Kevin Dalleau, Marie-Dominique Devignes, William Digan, Chia-Ju Lee, Ndeye-Coumba Ndiaye, Nadine Petitpain, Patrice Ringot, Malika Smaïl-Tabbone, Yannick Toussaint, Adrien Coulet

## Abstract

Pharmacogenomics (PGx) studies how individual gene variations impact drug response phenotypes, which makes knowledge related to PGx a key component towards precision medicine. A significant part of the state-of-the-art knowledge in PGx is accumulated in scientific publications, where it is hardly reusable to humans or software. Natural language processing techniques have been developed and employed for guiding experts curating this amount of knowledge. But existing works are limited by the absence of a high quality annotated corpus focusing on PGx domain. This absence restricts in particular the use of supervised machine learning. This article introduces PGxCorpus, a manually annotated corpus, designed to fill this gap and to enable the automatic extraction of PGx relationships from text. It comprises 945 sentences from 911 PubMed abstracts, annotated with PGx entities of interest (mainly gene variations, genes, drugs and phenotypes), and relationships between those. We present in this article the corpus itself, its construction and a baseline experiment that illustrates how it may be leveraged to synthesize and summarize PGx knowledge.

## Background & Summary

Pharmacogenomics (or PGx) studies how individual gene variations impact drug response phenotypes [48]. This is of particular interest for the implementation of precision medicine, i.e. a medicine tailoring treatments (e.g. chosen drugs and dosages) to every patient, in order to reduce the risk of adverse effects and optimize benefits. Indeed, examples of PGx knowledge have already translated into clinical guidelines and practices [3,11], recommending the consideration of individual genotypes when prescribing some particular drugs. For example, patients with the allele *57:01 of the HLA gene are at high risk to present a hypersensitivity reaction if treated with abacavir, an anti-retroviral, thus should be genotyped for this gene before prescription [31].

Many scientific publications are reporting the impact of gene variants on drug responses, and Medline size (30 million articles as of Sept. 2019) makes it hard for humans or machines to get a full understanding of the state of the art of this domain. NLP (Natural Language Processing) techniques have been consequently developed and employed to structure and synthesize PGx knowledge [7, 14]. Previous works mainly investigated rule-based approaches [5, 8, 37] and unsupervised learning [22, 35], because of the absence of an annotated corpus. Supervised learning has also been experimented [4, 25, 34, 38, 49], but without appropriate corpora, most studies build train and test sets on the basis of PharmGKB, the reference database for this domain [46]. Because it is manually curated, PharmGKB provides a high quality referential for such task. Annotations provided by PharmGKB (i.e. 2 associated entities and the identifier of the PubMed article in support) result from the consideration by human curators of various knowledge sources: article text; tables and figures; and curator’s own knowledge of the domain. Consequently PharmGKB annotations result from a high level process that can hardly be compared to an NLP-only approach. In particular, most NLP efforts are restricted to open-access texts only, without considering background knowledge. In this sense, system evaluated in comparison with PharmGKB are tested on how they may guide the curation process, but not on how they can capture what is stated in texts.

In domains close to PGx, corpora have been annotated with biomedical entities of interest for PGx, but never with the three PGx key entities, i.e. drugs, genomic variations and phenotypes, in the same portion of text, and never with relationships specific to PGx. Hahn *et al.* [14] made a panorama of corpora related to PGx and the most pertinents are presented and compared with PGxCorpus in the next section. In contrast with existing corpora PGxCorpus encompasses all three key entities of interest in PGx (i.e. drugs, genomic variations and phenotypes) both at the corpus level (the corpus encompasses the three) and at the sentence level (most sentences encompass the three). In addition, both entities and relationships are labeled using detailed hierarchies, allowing to capture pharmacogenomic relationships with a finest level of granularity. Finally, PGxCorpus is larger than other corpora: it is 2 to 3 times larger in term of annotated relationships to other corpora that include genomic entities. Despite the existence of reference resources, in particular PharmGKB, and of alternative to classical supervised learning such as weak supervision or active learning, we believe that a high quality training set remains an asset for a domain and that PGxCorpus will serve both the PGx and the Bio-NLP communities.

This manuscript presents first PGxCorpus itself; its construction, in Methods; and a baseline experiment, in Technical Validation.

## Description of the dataset: PGxCorpus

### Position in regards with existing corpora

In domains close to PGx, corpora have been annotated with biomedical entities, but only few of them include relationships (see Hahn *et al.* [14] for a panorama, plus [20, 26, 42, 51]). Closer corpora are related to pharmacovigilance or genetic traits, then focusing on drug–adverse response or SNP–phenotype associations. To our knowledge, no corpus includes annotations of all three PGx key entities, i.e. drugs, genomic variations and drug response phenotypes; and no corpus annotates PGx relationships between these entities. Developed for pharmacovigilance, **EU-ADR** [43] is a corpus composed of three disjoint subcorpora of 100 PubMed abstracts each. Sentences are annotated with pairs of entities either *drug-disease, drug-target* or *target-disease*. In the same vein, **ADE-EXT** [12] consists of 2,972 MEDLINE case reports, annotated with *drugs* and *conditions* (e.g. diseases, signs and symptoms) and their relationships. **SNPPhenA** [1] is a corpus of 360 PubMed abstracts, annotated with *single nucleotide polymorphisms* (SNPs), *phenotypes* and their relationships. Domains covered by EU-ADR, ADE-EXT or SNPPhena are related to PGx because they encompass entites of interest in PGx, but fit only partially the purpose of PGx relation extraction since sentences contains only two types of entities and that their annotated relations are only rarely concerning PGx. In particular, EU-ADR and ADE-EXT annotate drug reactions without considering genetic factors. SNPPhena does not focus on drug response phenotypes, but on general phenotype and symptoms, and it considers only SNPs whereas other genomic variations are of importance in PGx. In addition, the size of EU-ADR sub-corpora and SNPPhena are relatively small (only a few hundred annotated sentences), which limits the use of supervised learning approaches that require large train sets such as TreeLSTM [41]. These elements motivated us to construct a new corpus, focused on PGx, and large enough to train deep neural network models.

Table 1 proposes a comparison of PGxCorpus with five related corpora. In contrast with existing corpora and in particular those mentioned above, PGxCorpus encompasses all entities of interest in PGx (i.e. drugs, genomic variations and phenotypes) both at the corpus level and at the sentence level. Indeed, 47% of its annotated relations (1,353 out of 2,871) involve these three types of entities. The ratio of sentences with no relation in PGxCorpus is only 2.7%, which is rather low in comparison with larger corpora such as ADE-EXT where it is up to 71.1%. The number of types of relationships and entities employed in PGxCorpus are greater, allowing a finest level of granularity. Indeed, while PGxCorpus uses 10 types of entities types and 7 types of relations (all organized in hierarchies), SNPhenA, EUADE and ADE-EXT only use one type of relation (the relation occurs or not), but specify one of three modalities (positive, hypothetical or negative) in the case of EU-ADR and one of five modalities (neutral, weak confidence, moderate confidence, strong confidence and negative) in the case of SNPPhenA. In PGxCorpus, relationship modality (one out of the four values: positive, hypothetical, negative or both) is also captured and complements the relation type. SemEval DDI uses four types of drug-drug relations; however those are specific to drug-drug interactions and irrelevant to PGx. Unlike the other corpora, PGxCorpus includes nested and discontiguous entities, allowing to capture complex entities and their relationships. Finally, PGxCorpus is larger than other corpora that include genomic entities (required in PGx): it is 2 to 3 times larger in term of annotated relationships to SNPPhena and EU-ADR.

**Table 1.** Main characteristics of PGxCorpus in comparison with related corpora. Sizes of corpora are reported in term of number of sentences (sent.) and annotated relationships (rel.). Numbers of sentences without any annotated relation (Sent. w/o rel.) is reported both as a percentage (%) and an absolute number of sentences (nb.). The specific presence of PGx key entities, i.e. drugs (Dru.), genetic factors (Gen.) and phenotypes (Phe.) is reported under the Key entities column. Overall numbers of types of entities and relations used in annotations are reported as #Ent. and #Rel. types respectively. #Mod. refers to the number of modalities for the annotation of relations (e.g. positive, hypothetical, negative).

| Corpus name | Subcorpus | Corpus size |  | Sent. w/o rel. |  | Key entities |  |  | #Ent. types | #Rel. types | # Mod. | Nested entities | Discont. entities |
| --- | --- | --- | --- | --- | --- | --- | --- | --- | --- | --- | --- | --- | --- |
|  |  | sent. | rel. | % | nb. | Dru. | Gen. | Phe. |  |  |  |  |  |
| SNPPhenA | – | 483 | 1300 | 0 | 0 |  | ✓ | ✓ | 2 | 1 | 5 | no | no |
| EU-ADR | drug-disease | 244 | 176 | 0 | 0 | ✓ |  | ✓ | 2 | 1 | 3 | no | no |
|  | drug-target | 247 | 310 | 0 | 0 | ✓ | ✓ |  | 3 | 1 | 3 | no | no |
|  | target-disease | 355 | 262 | 0 | 0 |  | ✓ | ✓ | 3 | 1 | 3 | no | no |
| SemEval | DrugBank | 5,675 | 3,805 | 65.9 | 3739 | ✓ |  |  | 4 | 4 | 1 | no | no |
| 2013 DDI | MEDLINE | 1,301 | 232 | 87.1 | 1133 | ✓ |  |  | 4 | 4 | 1 | no | no |
| ADE-EXT | – | 5,939 | 6,701 | 28.9 | 1719 | ✓ |  | ✓ | 2 | 1 | 1 | no | no |
| **PGxCorpus** | – | 945 | 2,871 | 2.7 | 26 | ✓ | ✓ | ✓ | 10 | 7 | 4 | yes | yes |

Despite the existence of reference resources, in particular PharmGKB, and of alternative to classical supervised learning such as weak supervision or active learning, we believe that a high quality training set remains an asset for a domain and thus that the PGx community will benefit from PGxCorpus.

### Statistics on PGxCorpus

PGxCorpus encompasses 945 sentences, from 911 distinct PubMed abstracts, annotated with 6,761 PGx entities and 2,871 relationships between them. Detailed statistics on the type of entities and relationships annotated are provided in Table 2 and 3, respectively. Note that we distinguish two types of particular entities: nested and discontiguous ones. Nested entities are entities that encompass fully or partially at least one other entity in their offset. In Figure 1, the phenotype “acenocoumarol sensitivity" is an example of nested entity since it encompasses the “acenocoumarol” drug. Discontiguous entities are entities which offset is discontiguous, such as “VKORC1 genotypes” in Figure 1. 874 out of the 945 sentences of PGxCorpus (92%) contain the three key entities of pharmacogenomic (i.e., drug, genomic factor and phenotype). Figure 2 presents two additional examples of sentences of PGxCorpus. More examples can be browsed at https://pgxcorpus.loria.fr/. All the corpus abstracts were published between 1952 and 2017.

**Table 2.** Numbers of entities annotated in PGxCorpus, by type. Because nested and discontiguous (Discont.) entities are considered particularly in our baseline experiments, we report numbers of “simple” annotations, i.e. those that are neither nested nor discontiguous. Nested and Discont. refers to annotations that are either nested or discontiguous. “Both N&D” refers to entities both nested and discontiguous.

| PGxCorpus entity | Simple | Nested | Discont. | Both N&D | Total |
| --- | --- | --- | --- | --- | --- |
| Chemical | 1,512 | 192 | 2 | 12 | 1,718 |
| Genomic_factor | 21 | 68 | 7 | 3 | 99 |
| ↳ Gene_or_protein | 1,685 | 20 | 3 | 0 | 1,708 |
| ↳ Genomic_variation | 14 | 37 | 3 | 0 | 54 |
| ↳ Limited_variation | 237 | 537 | 98 | 47 | 919 |
| ↳ Haplotype | 15 | 112 | 4 | 6 | 137 |
| Phenotype | 282 | 330 | 60 | 27 | 699 |
| ↳ Disease | 460 | 143 | 14 | 18 | 635 |
| ↳ Pharmacodynamic_phenotype | 157 | 390 | 60 | 25 | 632 |
| ↳ Pharmacokinetic_phenotype | 31 | 109 | 14 | 6 | 160 |
| Total | 4,414 | 1,938 | 265 | 144 | **6,761** |

**Table 3.** Numbers of relations annotated in PGxCorpus, by type. Because of their relatively rareness, annotations made with ‘metabolizes’ or ‘transports’ types have been subsequently turned in as ‘influences’ annotations in the corpus. All counts are disjoint.

|  |  |
| --- | --- |
| isAssociatedWith | 733 |
| ↳ influences | 937 |
| ↳ causes | 168 |
| ↳ decreases | 263 |
| ↳ increases | 243 |
| ↳ treats | 238 |
| isEquivalentTo | 293 |
| Total | 2,875 |

**Figure 1.**
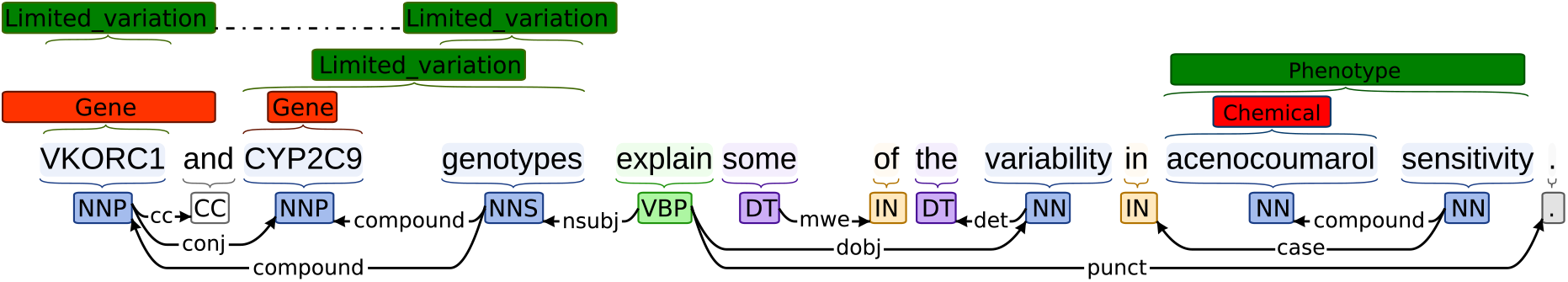
Example of sentence with PGx key and composite entities. The key entities, in red, correspond to entities retrieved by PubTator. Composite entities, in green, were obtained using the PHARE ontology. The syntactic dependency analysis is presented on the bottom of the figure and the entities on top.

**Figure 2.**
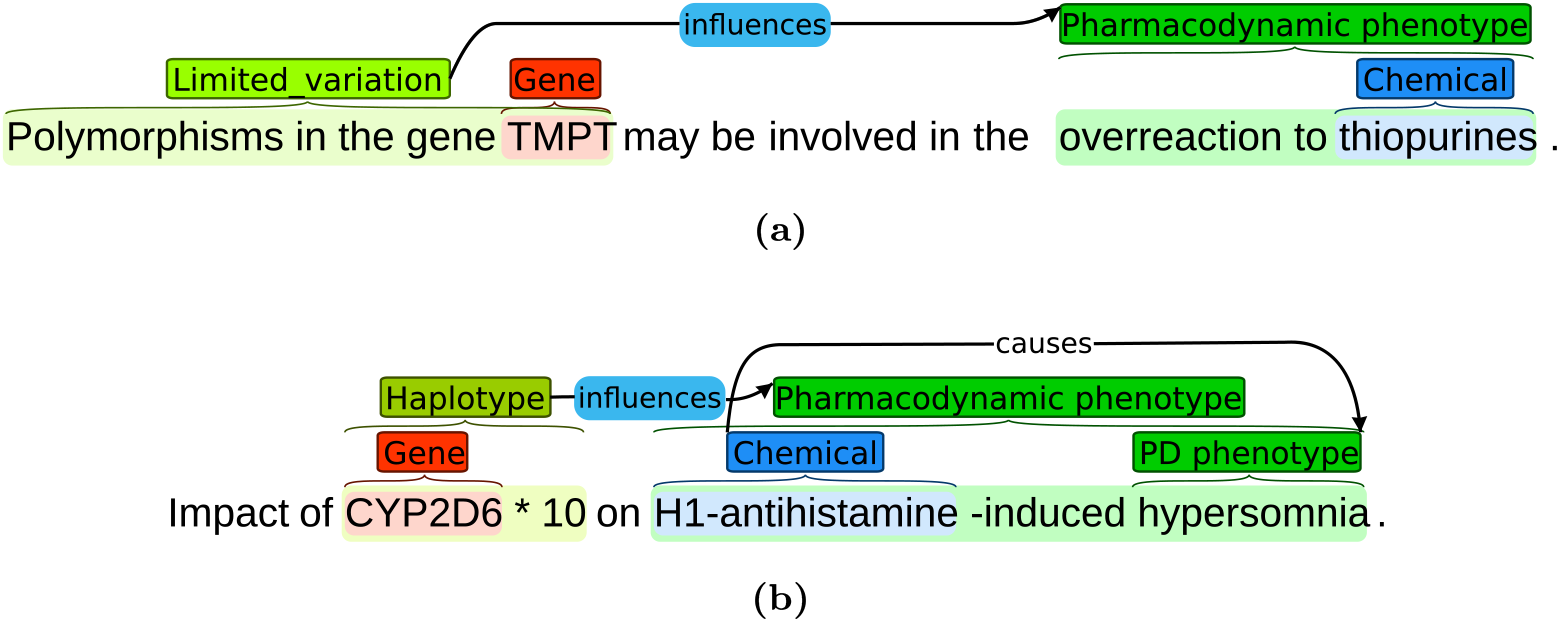
Two annotated sentences of PGxCorpus. Sentence *(a)* encompasses a relationship of type *influences* and of modality *hypothetical*, denoted by the blue color. Sentence *(b)* is a title, with two annotated relationships. The first is a relationship of type *influences* and of modality *hypothetical*. It is hypothetical because the title states that the paper studies the relation, but not that it is valid. The second relationship is of type *causes* and annotates a nominal group.

## Methods

In this section, we detail the steps of construction of PGxCorpus, as overviewed by Figure 3. This construction consists in two main steps: *(1)* the automatic pre-annotation of named entities and *(2)* the manual annotation that encompasses the correction of the pre-annotation and the addition of typed relationships between named entities.

**Figure 3.**
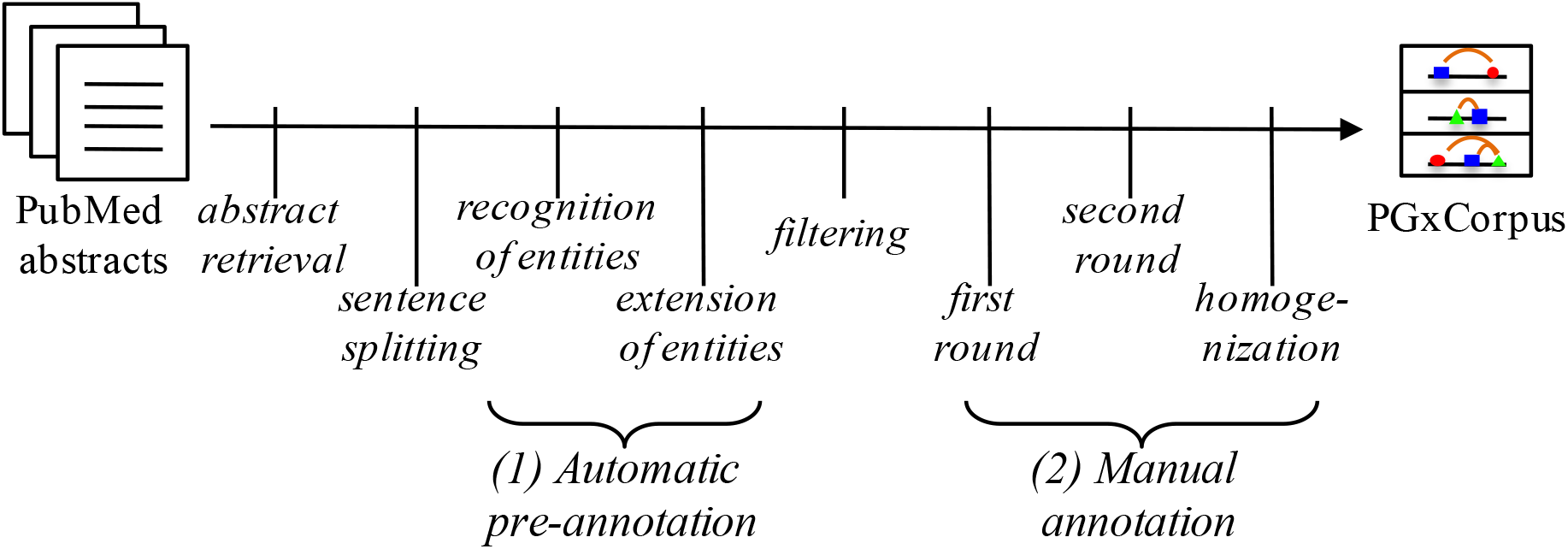
Overview of the construction of PGxCorpus.

We followed good practices proposed in [27], as well as practical examples provided by EU-ADR, ADE-EXT, SNPPhena and other corpora used in NLP shared tasks such as GENIA [21], SemEval DDI [15]. We particularly considered reports on the MERLOT corpus, which focuses on its annotation guidelines [2, 33] and inter-annotator agreement [10].

### Abstract retrieval and sentence splitting

The very first step consists in retrieving from PubMed [30] abstracts of publications related with PGx. This was performed with the tool EDirect [19] queried with:

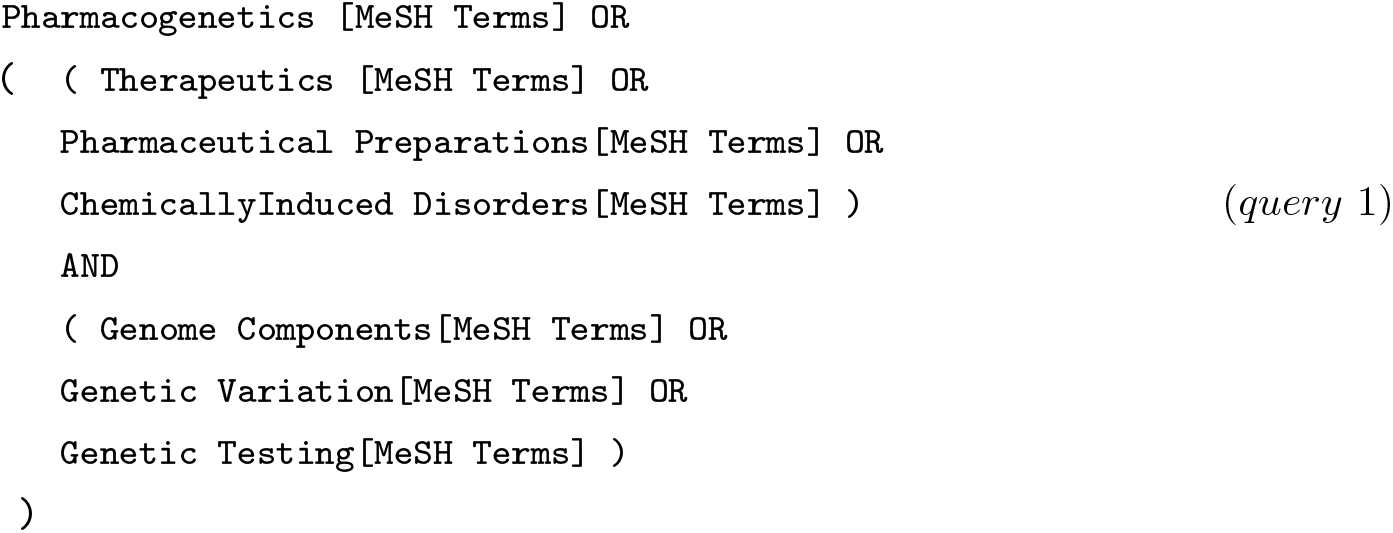

This query aims at retrieving article abstracts concerned with PGx or with at least one treatment and one genetic factor. It has been built by browsing manually the hierarchy of the MeSH vocabulary, which annotates PubMed entries. The use of MeSH terms allows PubMed to retrieve articles using synonyms and descendant terms of those used in the query. The query was voluntarily made general to retrieve a large set of abstracts that might mention PGx relationships.

Every retrieved abstract is subsequently split into its constitutive sentences, using GeniaSS [39].

### Automated pre-annotation

To facilitate the manual annotation of PGx relationships, we pre-annotate automatically sentences with various types of entities of interest for PGx. This pre-annotation is composed of two phases: First, PGx *key entities*, i.e. Gene, Mutation, Disease and Chemicals, are recognized and annotated with a state-of-the-art Named Entity Recognition (NER) tool. Second, these annotations are extended when they take part in the description of a PGx *composite entity*, such as a gene expression or a drug response phenotype.

#### Recognition of key PGx entities

Pre-annotation is initiated using PubTator [45], which recognizes the following biomedical entities from PubMed abstracts: chemicals, diseases, genes, mutations and species. PubTator integrates multiple challenge-winning text mining algorithms, listed in Table 4 along with their performances on various benchmark corpora. Disease recognition is performed with DNorm, which uses BANNER [23], a trainable system using Conditional Random Fields (CRF) and a rich feature set for disease recognition. For genes, GeneTUKit uses a combination of machine learning methods (including CRFs) and dictionary-based approaches. For mutations, tmVar also uses a CRF-based model with a set of features including dictionary, linguistic, character, semantic, case pattern and contextual features. PubTator was chosen for three reasons: it offers a wide coverage of PGx key entities; it provides an easy-to-use API to recover PubMed abstracts along with entity types and their boundaries; and it includes high performance NER tools.

**Table 4.** Performances reported for PubTator. PubTator is the NER tool used during the pre-annotation step of PGxCorpus. P, R and F1 stand for Precision, Recall and F1-score, respectively. n/a denotes we were not able to find information to fill the cell.

| Entity type | Tool | Evaluated on | Performance |  |  |
| --- | --- | --- | --- | --- | --- |
|  |  |  | P | R | F1 |
| Chemicals | Dictionary-based [47] | n/a | n/a | n/a | 53.82 |
| Disease | DNorm [24] | NCBI Disease Corpus | 82.8 | 81.9 | 80.9 |
| Gene | GeneTUKit [18] | n/a | n/a | n/a | 82.97 |
|  |  | GNAT-100 | 43.0 | 56.7 | 48.9 |
| Mutation | tmVar [44] | MutationFinder Corpus | 98.80 | 89.62 | 93.98 |

#### Extension of the annotations with the PHARE ontology

The second phase of the pre-annotation consists in extending automatically key entity annotations, when possible, with the PHARE (PHArmacogenomic RElationships) ontology [8]. This ontology encompasses frequent terms that, associated in nominal structure with PGx *key entities*, form PGx *composite entities*. These terms were obtained by analyzing dependency graphs of nominal structures in which a key entity syntactically *modifies* another term, and in turn were structured in the PHARE ontology. In the example provided in Figure 1, the drug name **acenocoumarol** syntactically modifies the term **sensitivity**. According to the PHARE ontology, the term *sensitivity*, when modified by a drug, forms a composite entity belonging to the *DrugSensitivity* class. Since this class is a subclass of the *Phenotype* class, **acenocoumarol sensitivity** may also be typed as a *Phenotype*. Following this principle, annotations of PGx key entities made by PubTator are extended, when possible, to PGx composite entities, then typed with classes of the PHARE ontology. For this matter, the dependency graph of each sentence is constructed with the Stanford Parser [9] and in each graph, the direct vicinity of key entities is explored in the search for terms defined in PHARE.

To homogenize the types of entities in PGxCorpus, we defined a reduced set of entities of interest, listed in Figure 4 and then defined mappings from PubTator entities and PHARE classes on one side to the types allowed in PGxCorpus on the other side. These mappings are reported in Table 5. Note that we decided to use a type Chemical, instead of Drug, first because we rely on PubTator that recognizes chemicals (without distinguishing between those and drugs), second because it allows to include broadly more candidate entities that may be involved in PGx relationships, such as drug metabolites or not yet approved drugs. Also, we decided on a type named Gene_or_protein, broader to Gene, because it is hard to disambiguate between gene and protein names in NLP, and commonly assumed that the task of gene name recognition is indeed a gene-or-protein name recognition [50].

**Table 5.** Mapping between PubTator entities types, PHARE classes and PGxCorpus entity types.

| Origin | Initial type | Type in PGxCorpus |
| --- | --- | --- |
| *PubTator* | Chemical | Chemical |
|  | Disease | Disease |
|  | Gene | Gene_or_protein |
|  | Mutation | Limited_variation |
| *PHARE* | Drug | Chemical |
|  | DrugMetabolite | Chemical |
|  | Gene | Gene_or_protein |
|  | GenomicRegion | Genomic_factor |
|  | GenomicVariation | Genomic_variation |
|  | GeneProduct | Gene_or_protein |
|  | Mutation | Limited_variation |
|  | Phenotype | Phenotype |

**Figure 4.**
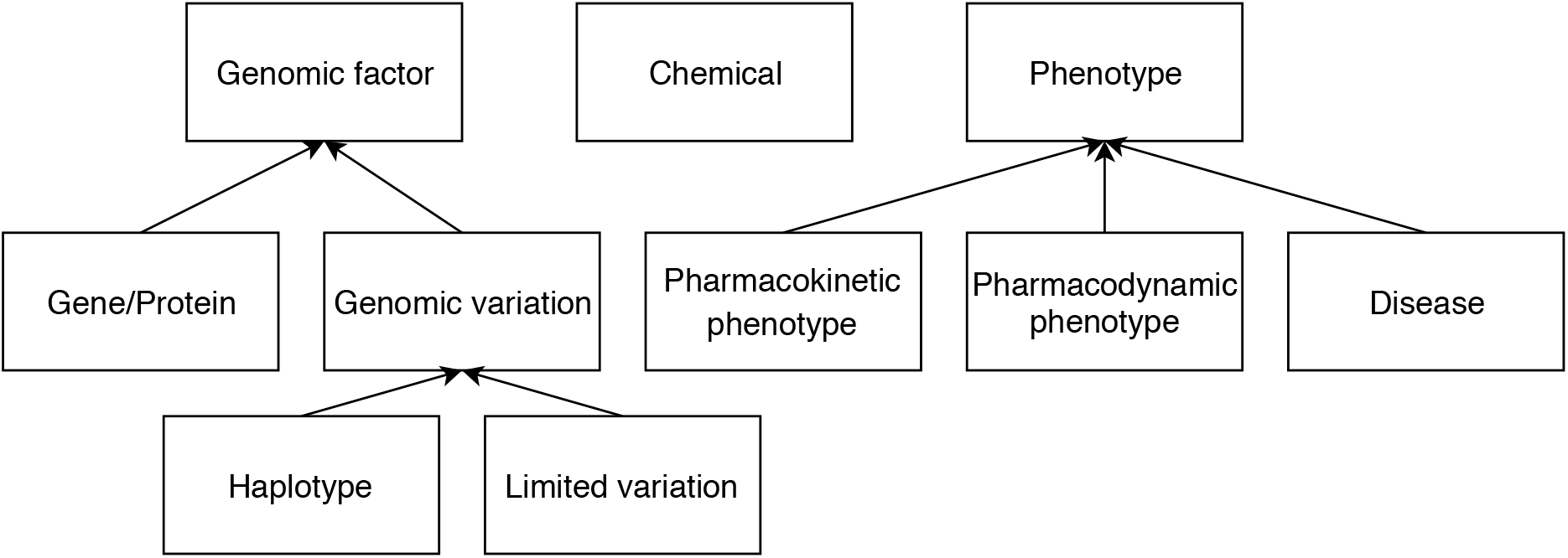
Types of entities annotated in PGxCorpus and their hierarchy.

### Manual annotations

Before the manual annotation itself, malformed sentences (sentence tokenization errors) and sentences that did not contain at least one drug and one genetic factor, according to PubTator or PHARE are filtered out.

Out of the remaining sentences, we randomly select 1,897 of them to be manually annotated. The annotation process is realized by 11 annotators, out of which 5 are considered senior annotators. Annotators are either pharmacists (3), biologists (3) or bioinformaticians (5). Each sentence is annotated in **three phases**: First, it is annotated independently by two annotators (senior or not); Second, their annotations are, in turn, compared and revised by a third, senior annotator; Last, a homogenization phase ends the process.

During the first phase, annotators are provided with sentences and entity pre-annotations. At this stage, they correct pre-annotations, add potential relationships between them, and discard sentences which are ambiguous or not related with PGx domain. Sentences discarded by at least one annotator are not considered for the second phase. During both first and second phases, sentences are randomly assigned to annotators, but we ensure that senior annotators revise only sentences they did not annotate in the first phase.

In order to ensure the consistency of the manual annotations, annotators are provided with **detailed guidelines** [29]. Those describe the type of entities and relationships to annotate (reported here in Figures 4 and 5), relationship modality (affirmed, negated, hypothetical), the main rules to follow, along with examples. Entity and relationship types are organized in simple hierarchies. Some of the relationship types are directly related to PGx (denoted with Δ in Figure 5), whereas some have a broader scope (i.e. isEquivalentTo and treats). Guidelines also provide an how-to-use guide for the annotation tool and answers frequently-asked questions. The first version of the guidelines was written before the first phase of the annotation. Examples and clarifications were regularly added to the document during the first phase of the annotation. Guidelines were subject to an important revision between the two first annotation phases, to clarify how to annotate ambiguous cases, which have been raised by annotators or by the evaluation of an agreement score between annotators (see Section Inter-annotator agreement).

**Figure 5.**
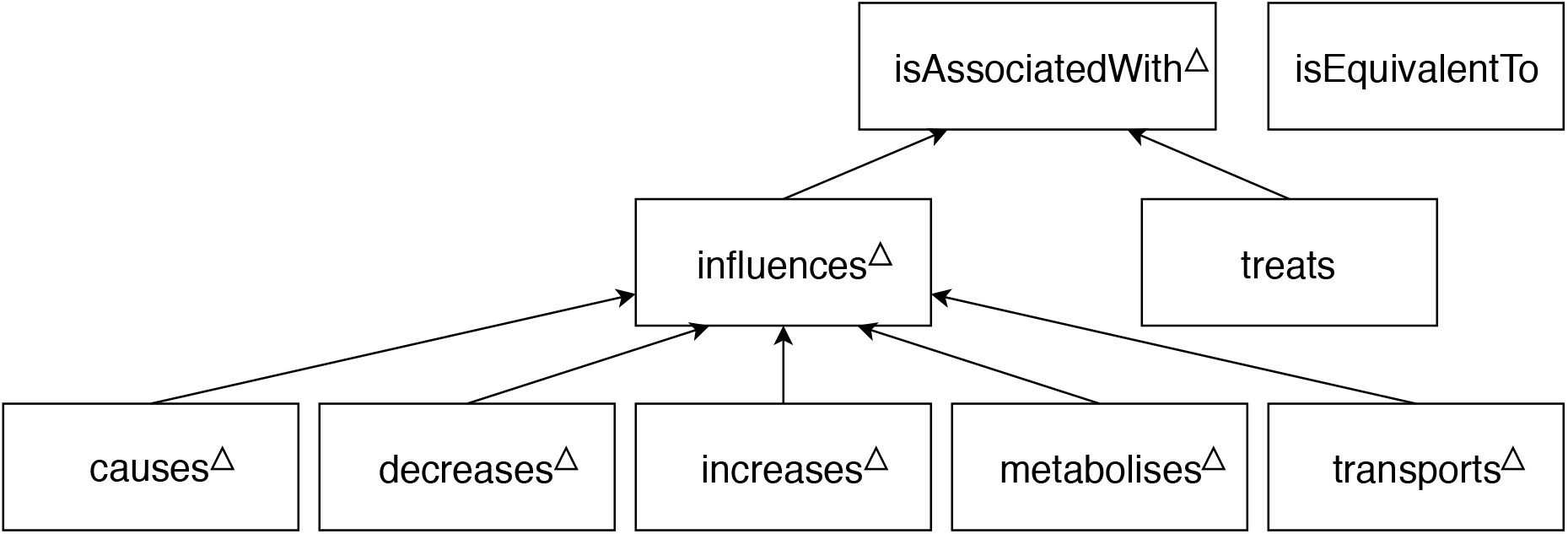
Types of relationships annotated in PGxCorpus and their hierarchy. Types directly related to PGx are marked with Δ, wheras isEquivalentTo and treats have a broader scope.

The final phase of **homogenization** ends the corpus construction process to reduce the heterogeneity that remains in the annotations after the second phase. Two expert annotators review together sentences in two times: the first time is a complete pass on all annotated sentences to identify sources of heterogeneity. The second time consists in *(a)* listing sentences associated with each source of heterogeneity using programmatic scripts and keywords, *(b)* reaching a consensus for their annotation, and *(c)* accordingly modifying the annotations. Sources of heterogeneity identified at this stage include: the annotation of drug combinations, of dose-related phenotypes, of mutation-related cancer types (e.g. p53-positive breast cancer), of behavior-related phenotypes (e.g. drug abuse, drug dependence), of genomic factors (e.g. exons, promoters, regulatory regions), of treated conditions (e.g. transplantations or post-surgery treatments), uncommon type of relationships. Concerning the latter, annotations made with uncommon types (i.e. ‘metabolizes’ and ‘transports’) are turned into their upper-level type of annotations (i.e. ‘influences’). This explains why ‘metabolizes’ and ‘transports’ types are in Figure 5, but not in Table 3. Regarding the cause of the heterogeneity in annotations, in some cases guidelines were specific but disregarded by annotators; in other cases they were caused by unexpected examples, absents from the guidelines.

### Code availability

A Git repository of the whole project is accessible at https://github.com/practikpharma/PGxCorpus/. It includes the annotation guidelines, the corpus itself and the programmatic code of the baseline experiments presented in Technical Validation.

## Data Records

### Data availability

PGxCorpus is available in the BioNLP shared task file format [36] at three locations:

- **figshare**, an open access data repository, at the following address: https://figshare.com/s/9d315cec6bb629d04210
- A **BRAT server** [40], enabling a friendly online visualization of the annotations: https://pgxcorpus.loria.fr/
- A **Git** repository of the whole project that also includes the annotation guidelines and programmatic code of the baseline experiments presented in Technical Validation https://github.com/practikpharma/PGxCorpus/.

### Statistics on the preparation of PGxCorpus

PubMed has been queried with our initial query (*query 1*) in July 2017, to retrieve 86,520 distinct abstracts, split out in 657,538 sentences. Statistics of pre-annotations obtained with PubTator and PHARE on these sentences are provided in Table 6 and 7, respectively. After filtering malformed sentences and sentences that do not contain at least one genomic factor and one drug, we obtain 176,704 sentences, out of which we randomly pick 1,897 sentences that are subsequently manually annotated. This number of sentences is chosen in regards of constraints of the distribution of the annotation task. These sentences come from 1,813 distinct abstracts.

**Table 6.** Type and number of entities recognized by PubTator in the pre-annotation.

| PubTator entity | Number<br>recognized |
| --- | --- |
| Chemical | 90,816 |
| Disease | 125,487 |
| Gene | 196,460 |
| Mutation | 25,417 |

**Table 7.** Number of entities pre-annotated after extending PubTator annotation with the PHARE ontology. Because discontiguous entities are excluded from our baseline experiments (see Section Technical Validation), their number is specified. No disease entity is annotated discontiguously first because PubTator is not generating discontiguous annotation, and second because the extension of annotations with PHARE (which may be discontiguous) is not producing disease annotations, but phenotype annotations.

| PHARE entity | Discontiguous | All |
| --- | --- | --- |
| Chemical | 430 | 87,764 |
| Disease | 0 | 29,589 |
| Gene_or_protein | 4,690 | 10,1326 |
| Genomic_variation | 8,698 | 13,601 |
| Phenotype | 10,935 | 16,770 |

The first phase of manual annotation, by 11 annotators, took roughly four months. The mean number of sentences annotated by an annotator is of 344.73 (standard deviation=126.33) sentences for this phase. The second phase, by 5 senior annotators, took four other months. Each senior annotator revised 258.6 (sd=0.54) sentences. Annotations were made on a voluntary basis, which explains the relatively long length of this process.

Sentences that were clearly not about PGx or presenting obvious problems such as incompleteness, typos or ambiguity were asked to be discarded. Accordingly, annotators discarded 952 sentences out of the 1,897 randomly picked, leaving 945 sentences. A very large majority of the motivation for discards was the scope of sentences. A typical example is the large number of sentences about genetic therapies annotators had to discard, since those also contain both a drug and a gene name and then were selected according to our filtering criteria.

## Technical Validation

In this section we present an inter-annotator agreement analysis and results of a baseline experiment of relation extraction where PGxCorpus is used as the training set of a neural network model.

### Inter-annotator agreement

#### Metrics

The annotation task considered for PGxCorpus is particularly rich: it employs 10 entity types, 9 relation types and 3 relation modalities (sometimes named attributes); in addition, entities may be nested or discontiguous. Given this richness, metrics to control the variability of the annotations have been evaluated, in particular at the end of the first phase of the manual annotation, when each sentence has been annotated independently by two annotators. We compute an agreement score that evaluates how much annotators agreed with each others using the F1-score, following [13, 17]. In this case, the agreement or F1-score, is measured using one annotator as a reference and the other as a prediction. Note that inverting the reference and the prediction only inverts the precision and the recall but has no effect on the F1-score itself. We preferred the F1-score instead of other conventional measures such as the kappa coefficient [6] because of the complexity of our annotation task. Kappa coefficient is designed to evaluate inter-annotator agreements while taking into account the probability that the agreement might be due to random guess. It is adapted when annotators select a category, out of a set, to annotate already marked-up entities. Then, larger the set is, the less probable an agreement occurs by chance. In our case, the annotators need not only to select a category, but also to identify the boundaries of these potential entities. In this setting, the probability of a by-chance agreement within the kappa coefficient is low and unadapted. The F1-score is defined as the harmonic mean of the precision 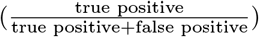 and recall 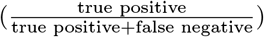, i.e. 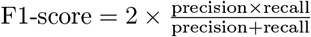.

#### Entity agreement

Agreement on entity annotations is determined in four ways, in regards with two parameters: *(a)* using *exact* or *partial* match; *(b)* considering the entity hierarchy or not.

a. An *exact match* occurs when two annotators agree on both the entity type and their boundaries. A *partial* match is more flexible since it occurs when two annotators agree on the entity type, but annotation boundaries only overlap. Note that an annotation from the first annotator may overlap with multiple annotations from the second annotator, and vice versa. Considering every overlapping entities as a match would artificially increase the recall and the precision because only one can indeed reflect an agreement between the two annotators. We ensure in this case that an entity from the first annotator is matched with at most one entity from the second annotator using the Hopcroft-Karp algorithm [16]. In this case, the problem is seen as a maximum matching problem in a bipartite graph, where each set of annotations, one for each annotator, represents a sub-graph. The edges between the two sets represent possible overlaps between one annotation from the first annotator and one from the second.
b. We also consider a more flexible setting where the agreement takes into account upper hierarchies of entities and relationships, as defined in Figures 4 and 5. For instance, if a first annotator annotates an entity as *Pharmacokinetic phenotype (PK)* and a second as *Pharmacodynamic phenotype (PD)*, we consider they agreed to some extent, since both are subtype of *Phenotype*. In this setting, it can be considered that an entity (or relationship) is indeed annotated with several types: the one specified by an annotator and its parents in the hierarchy. In practice, if we consider the first annotator as the reference and the second as the prediction, we can distinguish three cases: (1) the prediction is more specific than the reference. In this case, common annotations shared by reference and prediction are counted as *true positives*, while annotations of the prediction that are too specific are *false positives*. For instance if the reference is *Phenotype* and the prediction is *PD*; we count one *false positive* in the evaluation of *PD* predictions, but the additional *Phenotype* annotation, inferred from the hierarchy, enables to count one *true positive* for *Phenotype* predictions. (2) The prediction is less specific than the reference. In this case, common annotations shared by reference and prediction are counted as *true positives*, while classes from the reference that are missed by the prediction are *false negative*. For instance if the reference is *PD* and the prediction is *Phenotype*, we count one *true positive* for *Phenotype* prediction, but one *false negative* in the prediction of *PD*. (3) The reference and the prediction do not have a direct hierarchy relationships, but a common ancestor (like *PD* and *PK*). In this case classes that are shared by the prediction and reference (i.e. the common ancestors) are counted as *true positive*, but too specific predictions as *false positives* and missed predictions as *false negatives*. For instance if the reference is *PD* and the prediction is *PK*, we count one *true positive* for the prediction of *Phenotype* (i.e. the only common ancestor), one *false positive* for the prediction of *PK* and one *false negative* for the prediction of *PD*.

Table 8 presents the inter-annotator entity agreement scores, obtained after the first phase of manual annotation, depending on settings *(a)* and *(b)*. We observe that for relatively simple entities such as chemicals, genes, haplotypes or diseases, F1-score even on the strictest constraints (exact match, no hierarchy) overpasses 70. We observe also that for more complex entities such as phenotype or genomic variations, annotators tend to agree on the presence of an entity, but not on its offset. This lead to some very low agreement. This motivates us to update the annotation guidelines between the two annotation phases, to clarify on how to decide on entity offsets. For instance it was clarified that adjective qualifying phenotypes should not be annotated, except if they are part of the name of the disease or symptom, but not simply qualifying its localisation or gravity. Accordingly when encountering “acute pain” annotators have to not annotate *acute*, but *pain*; whereas encountering “Acute Myeloid Leukemia”, a specific family of leukemia, annotators have to annotate all three words. When considering the hierarchy, the performances for the leaves of the hierarchy should not be affected. However, a slight drop is observed due to the use of the Hopcroft-Karp algorithm. Indeed, when using the hierarchy more potential matches can be observed between prediction and reference annotations generating more edges in the associated bipartite graph. The Hopcrof-Karp algorithm then removes some of the correct matches between leaves, causing a slight drop in the recall.

**Table 8.** Inter-annotator agreement (F1-score) for entity annotations. Four different settings enabling more or less flexibility are presented. The agreement score is computed after the first phase of manual annotation.

| **Entity matching:** (exact or partial) | exact | exact | partial | partial |
| --- | --- | --- | --- | --- |
| **Considering hierarchy:** (yes or no) | no | yes | no | yes |
| Chemical | 76.8 | 76.8 | 82.1 | 82.1 |
| Genomic_factor | 38.6 | 72.6 | 38.8 | 85.7 |
| ↳ Gene_or_protein | 85.3 | 85.3 | 90.0 | 89.4 |
| ↳ Genomic_variation | 32.9 | 49.3 | 53.0 | 76.8 |
| ↳ Limited_variation | 50.8 | 50.8 | 69.0 | 66.2 |
| ↳ Haplotype | 76.2 | 76.2 | 77.2 | 76.1 |
| Phenotype | 30.5 | 51.0 | 53.9 | 72.6 |
| ↳ Disease | 71.3 | 71.0 | 80.9 | 79.1 |
| ↳ Pharmacokinetic_phenotype | 48.2 | 48.2 | 57.0 | 57.0 |
| ↳ Pharmacodynamic_phenotype | 31.7 | 31.7 | 47.0 | 47.0 |
| Macro average | 57.4 | 63.8 | 68.7 | 76.1 |

#### Relation agreement

Regarding the inter-annotator agreement on relation annotations, we consider the same two settings, plus an additional one: *(a)* using *exact* or *partial* match, which applies in this case to the two entities involved in the relation; *(b)* the consideration of the hierarchy, which applies in this case to both the hierarchy of entities and relations (see Figure 4 and 5); *(c)* the direction of the relation is considered or not. Resulting agreements are presented in Table 9.

**Table 9.** Inter-annotator agreement (F1-score) for the annotation of relations. Five different settings are presented.

| **Entity matching:** | exact | exact | partial | partial | partial |
| --- | --- | --- | --- | --- | --- |
| **Considering hierarchies:** | none | both | none | both | both |
| **Considering direction:** | yes | yes | yes | yes | no |
| isAssociatedWith | 12.6 | 14.3 | 13.2 | 33.3 | 33.3 |
| ↳ influences | 12.8 | 12.8 | 17.7 | 29.3 | 29.8 |
| ↳ causes | 35.8 | 35.2 | 37.6 | 37.2 | 39.6 |
| ↳ decreases | 25.8 | 26.8 | 33.6 | 36.7 | 36.7 |
| ↳ increases | 14.5 | 15.6 | 27.4 | 30.2 | 30.2 |
| ↳ metabolizes | 59.0 | 59.0 | 61.5 | 61.5 | 61.5 |
| ↳ transports | 83.1 | 83.1 | 83.1 | 83.1 | 83.1 |
| ↳ treats | 33.2 | 34.7 | 36.3 | 37.3 | 37.3 |
| isEquivalentTo | 39.6 | 40.2 | 40.7 | 41.3 | 62.5 |
| Macro average | 47.3 | 47.1 | 50.3 | 53.8 | 57.0 |

Although the agreement on the relations is low, note that a relation can be considered correct only if an initial agreement on the two entities in relation has been reached.

### Pre-annotation correction

To evaluate how much annotators had to correct the automatically computed pre-annotations, we measured an agreement score between automatic pre-annotations and final annotations. As for the interannotator agreement, this is computed in the form of an F1-score. In this case, the final annotation is considered as the gold standard, and the F1-score measures how automatic pre-annotation performs in regards with the final annotation ground truth. Table 10 presents F1-score agreements in four distinct settings, enabling to consider how much the boundaries or the type of pre-annotations (or both) needed corrections. The precision of 94.1% when boundaries and type are relaxed means that pre-annotations point at offsets where something is indeed to annotate (94.1% of the time). The relatively low recall in the relaxed setting (64%) illustrates the amount of fully new annotations added by annotators. When considering stricter setting, with an exact match or a same type or both we observe that precision is getting lower, illustrating that entity boundaries and types frequently require corrections. Note that this agreement is relevant only for named entity, but not for relations because the latter are not pre-annotated, but fully done manually.

**Table 10.** Agreement (F1-score) between pre- and final annotations. This agreement evaluates the amount of manual corrections that was required after the automatic pre-annotation phase. Numbers in parenthesis provide the balance between precision (P) and recall (R).

| **Entity matching:** | exact | exact | partial | partial |
| --- | --- | --- | --- | --- |
| **Type matching:** | same type | potentially different | same type | potentially different |
| F1-Score (P;R) | 54.2 (66.9;45.5) | 59.0 (72.9;49.6) | 64.4 (79.6;54.1) | 76.2 (94.1;64.0) |

### Baseline experiments

In this section, we report on baseline experiments with PGxCorpus, which evaluates quantitatively its usefulness for extracting PGx entities and relations from text. The task evaluated here is composed of a first step of named entity recognition (NER) and a second one of relation extraction (RE). The NER is achieved with a variant of a Convolutional Neural Network (CNN) model, whereas the RE is processed with a multichannel CNN (MCCNN). Both models are detailed in the supplementary methods section of the supplementary material. Source code of the experiments is available at https://github.com/practikpharma/PGxCorpus/.

In a related work [32], we used a preliminary, partial and naive set of annotations, for testing the feasibility of extracting relations and incorporating them in a knowledge network. This included only 307 sentences (out of 945), annotated with a simplified schema of only 4 entity types and 2 relation types. The associated model for RE was simplistic, since it aimed at proofing feasibility only. The baseline experiment reported here considers all sentences of PGxCorpus and has been done with more advanced annotation schema and models.

#### Baseline performances

The objective of these experiments was not to reach the best performances but rather to propose a baseline for future comparisons, as well as to empirically demonstrate the usefulness of PGxCorpus for extracting PGx entities and relations from text.

##### Named entity recognition

Performances for the named entity recognition experiments, evaluated with a 10-fold cross validation, are reported in Table 11. A main limitation of the NER model is that discontiguous entities were not considered. This may hurt the performance even for contiguous entities since discontiguous entities were considered as negative, even though they might be very similar (from the model point of view) to contiguous entities.

**Table 11.** Performances of the task of named entity recognition in terms of F1-score (and its standard deviation in brackets, for the last setting). Balance between precision and recall, as well as details on standard deviations are provided in Supplementary Table S1.

| Entity matching: (exact or partial) | exact | exact | partial | partial |
| --- | --- | --- | --- | --- |
| Considering hierarchy: (yes or no) | no | yes | no | yes |
| Chemical | 76.07 | 76.07 | 82.67 | 82.67 (7.24) |
| Genomic_factor | 22.86 | 71.41 | 27.68 | 83.19 (5.90) |
| ↳ Gene_or_protein | 85.72 | 85.72 | 90.58 | 90.05 (3.89) |
| ↳ Genomic_variation | 2.67 | 49.13 | 3.83 | 71.18 (9.55) |
| ↳ Limited_variation | 47.08 | 47.02 | 72.71 | 71.57 (9.50) |
| ↳ Haplotype | 66.97 | 66.97 | 72.47 | 72.47 (19.34) |
| Phenotype | 31.76 | 50.80 | 48.48 | 69.57 (5.40) |
| ↳ Disease | 66.90 | 66.88 | 75.68 | 72.59 (7.30) |
| ↳ Pharmacokinetic_phenotype | 29.30 | 29.30 | 36.47 | 36.27 (19.40) |
| ↳ Pharmacodynamic_phenotype | 38.54 | 38.50 | 58.84 | 58.18 (10.11) |
| Macro average | 49.15 | 59.11 | 59.76 | 71.93 (5.64) |

From results reported in Table 11, several observations can be made. First, the best performances were obtained for *Chemical*, *Gene_or_protein* and *Disease* types, for which (1) the number of training samples is high, (2) PubTator annotations are available and (3) the ratio between normal entities and nested and/or discontiguous entities is low (see Table 2). Note that the definition for the *Limited_variation* entity used in our corpus is broader than the *Mutations* recognized by PubTator. PubTator recognizes precises descriptions of variations such as “VKORC1:C>A”, but not general ones such as “a VKORC1 polymorphism”, which we consider. This explains why the performances obtained for *Limited_variation* were lower than those obtained with PubTator (see Table 4). Even though the number of training samples for *Pharmacokinetic_phenotype* and *Haplotype* is low, we obtained reasonable performances. This may be due to a rather homogeneous phrasing and syntax in the mention of these entities. Not considering the hierarchy in cases as *Genomic_variation* or *Genomic_factor* types for which few training samples are available and a high heterogeneity is observed led to poor performances. Lastly we note that, as expected, the standard deviation for classes with only few examples annotated was high or very high (above 19 for *Haplotype* and *Pharmacokinetic_phenotype*). The random distribution of these “rare” examples between train and test sets, in the 10-fold cross validation, had a strong impact on performances, and explains large standard deviations. Concerning concepts that are leaves of the hierarchy, we observed a slight drop in performances when considering the hierarchy. This is due to the use of the Hopcroft-Karp algorithm as mentioned in the Entity agreement Subsection.

##### Relation extraction

Performances for the relation extraction (RE) experiments, evaluated with a 10-fold cross validation, are reported in Table 12. The RE model faced several limitations. (1) For a given sentence along with identified entities, the relation predictions were independent. This is obviously too simplistic and the prediction should be made globally. (2) We considered relationships annotated as *negated* or *hypothetical* by annotators just as regular relationships.

**Table 12.**
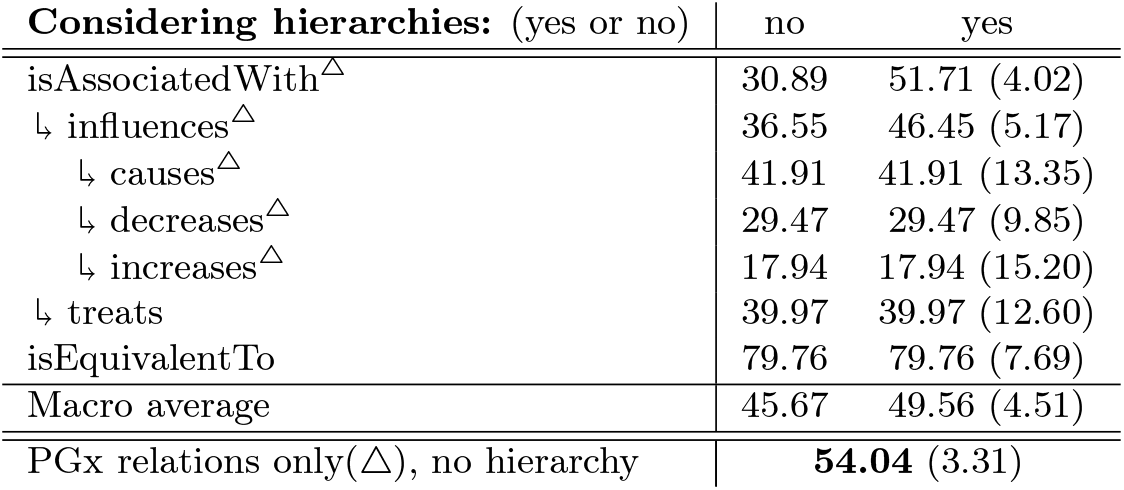
Performances of the task of relation extraction in terms of F1-score (and standard deviation). The last line provides results of an experiment for which only one category is considered, merging all the types specific to PGx (marked with Δ). For leaves, performances are unchanged when considering the hierarchy. Balance between precision and recall, as well as details on standard deviations are provided in Supplementary Table S2.

Several observations can be made about the RE results reported in Table 12. First, the fact that the model had to deal with multiple, complex and associated classes made the classification problem difficult and the performances relatively modest. The experiment in which we considered the hierarchy showed that, even if it was difficult to identify a specific type of relation, is was easier for the model to determine whether there was a relation between two entities or not. In other words, many mis-classifications were in fact predictions for types that belong to the same branch of the hierarchy. Like for the NER, types of relations with less examples tended to be associated with poorer performances and higher standard deviations (except for the *isEquivalentTo* relationship, which is very homogeneous). To build upon these observations, and particularly to avoid the impact of isEquivalentTo type that is not specific to PGx, we evaluated how PGxCorpus can be used to train a model for any relation specific to PGx (denoted with Δ in Table 12), without considering sub-types and the hierarchy. Results of this experiment is provided on the last line of Table 12

Several enhancements could be proposed to improve this baseline model. First, in our implementation, the hierarchy was not considered during the training phase. Accordingly, learning to predict a leaf penalized all the other categories, even those that were considered correct at test time. This explains why the “PGx Relations only” experiment led to better performances than individual classifications with or without hierarchy. On the other hand, considering the hierarchy at training would increase the number of examples for the higher categories of the hierarchy, potentially harming performances for the leaves. A model enabling multiclass labeling and a weighting dependent on the size of classes should balance this bias.

### Building upon PGxCorpus

We assembled a manually annotated corpus for pharmacogenomics, named PGxCorpus, and provide an experimental validation of its usefulness for the tasks of NER and RE in this domain.

Unlike existing corpora, PGxCorpus encompasses all three key entities involved in PGx relationships (drugs, genomic factors and phenotypes) and provides a fine-grained hierarchical classification for both PGx entities and relationships. By making this corpus freely available, our objective is to enable the training of supervised PGx relation extraction systems and to facilitate the comparison of their performances. Furthermore, the baseline experiment illustrates that PGxCorpus enables studying many challenges inherent with biomedical entities and relationships such as discontiguous entities, nested entites, multilabeled relationships, heterogenous distributions, *etc.*. In particular, PGxCorpus offers both a training resource for supervised approaches and a reference to evaluate and compare to future efforts. Out of PGx, such a corpus may serve connected domains by the use of transfer learning approaches, as illustrated by [28]. For all these reasons, we think that tasks of PGx NER and RE, supported by the novel existence of PGxCorpus, are well suited for proposing NLP Challenges and shared tasks. By this mean we expect that PGxCorpus will stimulate NLP research as well as facilitating the synthesis of PGx knowledge.

## Usage Notes

PGxCorpus is made available under the Creative Commons Attribution-Non-Commercial 4.0 International Public License. The programmatic code of our baseline experiments is available at https://github.com/practikpharma/PGxCorpus/tree/master/baseline_experiment.

## Acknowledgements

The authors acknowledge their funding institutions: the French National Research Agency (ANR) supports the PractiKPharma project (ANR-15-CE23-0028) and the University of Lorraine program “Lorraine Université d’Excellence” (15-IDEX-0004); Inria supports the *Snowball* Inria Associate Team.

## Author contributions

JL conducted the annotation campaign, designed and conducted baseline experiments, and wrote the manuscript.

RG conducted the annotation campaign.

AC, CB, CJL, JL, KD, MDD, MST, NCN, NP, RG, WD annotated the corpus and reviewed the manuscript.

PR advised on technical aspects of the project and set up the annotation servers.

YT advised on the design of the project and in the writing of the manuscript.

AC designed the study, supervised the annotation campaign and wrote the manuscript.

All authors read and approved the final manuscript.

