## Supplementary information for "PGxCorpus: a Manually Annotated Corpus for Pharmacogenomics"

### PGxCorpus – Supplementary information

This supplementary document includes:

- Supplementary methods about the representation and models used for baseline experiments.
- Table S1: Detailed performances of the task of named entity recognition.
- Table S2: Detailed performances of the task of relation extraction.

#### Supplementary methods

##### Sentence representation with word embeddings

Both our models for NER and RE are fed with *word embeddings* (*i.e.*, continuous vectors) of dimension $d_w$, along with extra *entity embeddings* of size $d_e$. RE is fed with an additional *nested entity embeddings* of size $d_n$.

Regarding word embeddings, given a sentence of $N$ words, $w_1, w_2, \ldots, w_N$, each word $w_i \in \mathcal{W}$ is embedded in a $d_w$-dimensional vector space by applying a lookup-table operation: $LT_W(w_i) = W_{w_i}$, where the matrix $W \in R^{d_w \times |\mathcal{W}|}$ represents the parameters to be trained in this lookup-table layer. The dictionary $\mathcal{W}$ is composed of all the words of the corpus. Each column $W_{w_i} \in R^{d_w}$ corresponds to the embedding vector of the $w_i$ word in our dictionary $\mathcal{W}$.

Beside word embeddings, two additional embeddings, named entity embeddings, are used to feed our models. (1) One entity embeddings enables to represent what type of entity a word composes. (2) One represents if the word starts, continues or ends the description of an entity. Both use a standard encoding of tags with Begin Intermediate Other End and Single (BIOES)-prefixes [6]. These two first entity embeddings are constructed slightly differently for NER and RE, since in the first, it encompasses tags for entities pre-annotated with PubTator and tags for entities annotated with PGxCorpus types, whereas in the latter, it considers tags for entity types of the corpus, plus special tags that marks pairs of entities between which a relationship may stand.

For the RE model only, a *nested entity embedding* of size $d_n$ is added to word and entity embeddings to represent entity types that may be included in nested entities involved in relations. For each word a *nested entity embedding* is added for each entity type. Given an entity type, this embedding can take one of two values: (a) *absent* if the word is not part of one of the two entities potentially related, or if it is part of one, but no entity of the given type is included in the entity of interest; (b) *present* if the word is part of one of the 2 entities and this one includes another entity of the given type.

Finally, word, entity and nested entity embeddings are concatenated to form the input corresponding to a given word. Let's denote $x_i$ the concatenated input corresponding to the $i^{th}$ word.

##### Named entity recognition model

The core of the CNN model used for NER is described in [2]. We adapted it, along with experiment settings, to fit with the particularity of PGxCorpus that is to encompass about one third of *discontiguous* or *nested entities* (2,347 discontiguous or nested / 6,761 entities, see Table 2).

Recognizing discontiguous entities is a complex and open problem in NLP and this baseline experiment does not aim at tackling it. For this reason, we discarded in the sentences, annotations of discontiguous entities from both our train and test sets (265/ 6,761 entities). Nested entities are considered in our experiment by applying the NER model recursively, as many times as there are nesting levels. Entities discovered during one iteration of the model are considered as input of the next iteration. Given the example of Figure 1, a first iteration will recognize the three entities "VKORC1", "CYP2C9" and "acenocoumarol". Then, the second iteration will consider them as an input to recognize "CYP2C9 genotypes" and "acenocoumarol sensitivity". "VKORC1 genotypes" is discontiguous and consequently discarded from the experiment.

Formally, given an input sequence $x_1, \ldots, x_N$, a classical sliding window approach is followed by applying a two-layer neural network (NN) on each possible window of size $k$. We denote $\mathcal{P}$ the set of BIOES-prefixed tags. Given the $i^{th}$ window, the NN computes a vector of scores $s_i = [s^1, \ldots, s^{|\mathcal{P}|}]$, where $s^t$ is the score of the BIOES-prefixed tag $t \in \mathcal{P}$, associated with the input $x_i$. Scores of the window $i$ are given by the following formula:

$$s_i = W_1 \; h( \; W_2 \; [x_{i-(\frac{k-1}{2})}, \ldots, x_i, \ldots, x_{i-(\frac{k+1}{2})}] \; ),$$

where the matrices $W_1 \in R^{d_h \times k|\mathcal{W}|}$ and $W_2 \in R^{|\mathcal{P}| \times d_h}$ are the trained parameters of the NN, and $h$ is a pointwise non-linear function such as the hyperbolic tangent, $d_h$ is the number of hidden units and

$k$ the size of the window. Inputs with indices exceeding the input boundaries, i.e. when $i - (\frac{k-1}{2}) < 1$ or $i - (\frac{k+1}{2}) > N$, are mapped to a special padding vector, which is also learned.

Scores of each window are finally given to a lattice module that allows to aggregate the BIOES-prefixed tags from our tagger module in a coherent manner, to recover the predicted labels. For more details about this layer, please see [2].

##### Relation extraction model

The model used for RE is a multichannel CNN (MCCNN) described in [5], where it has been successfully applied to the task of extraction of drug-drug and protein-protein interactions. It takes an input sentence and two recognized entities, computes a fixed size representation by composing input word embeddings. This representation is given to a scorer, which computes a score for each possible type of relationships. Sentences with more than two entities are considered by the model iteratively for each possible pair of entities for which a relation may stand, in both directions since relations may be oriented.

The MCCNN applies a CNN of variable kernel size to each input channels of word embeddings. In other words, it considers different embedding channels i.e. different versions of the word embeddings associated with each word, allowing to capture different aspects of input words. Formally, given an input sequence of word representations (i.e. concatenation of word and entity embedding) $x_1, \ldots, x_N$, applying a kernel to the $i^{th}$ window of size $k$ is done using the following formula:

$$C_i = h\Big(\sum_{j=1}^{N-k+1} W[x_i, \ldots, x_{i+k-1}]^j + b\Big)$$

where $[.]^j$ denotes the concatenation of inputs from channel $j$, $W \in \mathcal{R}^{(d_w+d_e)\times d_h}$ and $b \in \mathcal{R}^{d_h}$ are the parameters, $d_h$ is the size of the hidden layer, $h$ is a pointwise non-linear function such as the hyperbolic tangent and $N - k + 1$ is the number of input channels. For each kernel, a fixed size representation $r^* \in \mathcal{R}^{d_h}$ is then obtained by applying a max-pooling over time (here, the "time" means the position in the sentence):

$$r^* = \max\left[C_1, \ldots, C_{N-k+1}\right] .$$

We denote $K$ the number of kernels with different sizes. A sentence representation $r \in \mathcal{R}^{d_s}$ (with $d_s = K * d_h$) is finally obtained by concatenating the output corresponding to the $K$ kernels $r = [r_1^*, \ldots, r_K^*]$ .

The sentence representation is finally passed to a single layer NN, which outputs a score for each possible relation type:

$$s(r) = W^{(s)} r + b^{(s)} \ ,$$

where $W^{(s)} \in \mathcal{R}^{d_s \times |S|}$ and $b^{(s)} \in \mathcal{R}^{|S|}$ are the trained parameters of the scorer, $|S|$ is the number of possible relation types. The scores are interpreted as probabilities using a softmax layer [1].

##### Experimental settings

Word embeddings were pre-trained using the method described in [4] on about 3.4 million PubMed abstracts, corresponding to articles published between Jan. 1, 2014 and Dec. 31, 2016. Our models were trained by minimizing the negative log-likelihood over the training data. All parameters –embeddings, weights $W$ and biases $b$– were iteratively updated via backpropagation. We used a *hard tanh* function as activation function $f$. Hyper-parameters were tuned using a 10-fold cross-validation by selecting the values leading to the best averaged performance, and fixed for the rest of the experiment.

For NER, the CNN was fed with word embeddings and two types of entity embeddings (one with PubTator tags, used only for the first iteration of the model and one with PGxCorpus tags used in next iterations) of size $d_w = 100$ and $d_e = 20 \times 2$ (20 for each type of tags), respectively. The size of the hidden layer was fixed to $d_h = 200$, the kernel size to $k = 5$ and the learning rate to 0.01.

For RE, the MCCNN was fed with word embeddings and two types of entity embeddings (one with PGxCorpus entity tags; one to identify pairs of entities between which a relation may stand) of size $d_w = 200$ and $d_e = 20 \times 2$, respectively. The size of the nested entity embeddings was set to $d_n = 5 \times |\mathcal{E}|$, where $\mathcal{E}$ is the entity type dictionary.

We used two kernels of size 3 and 5. Following [3], both channels were initialized with pre-trained word embeddings, but gradients were backpropagated only through one of the channels. The size of the hidden layer was fixed to $d_h = 200$ and the learning rate to 0.01.

For both NER and RE, we applied a dropout regularization after the embedding layers [7] with a dropout probability fixed to 0.5. Both models were evaluated using a 10-fold cross validation. Each result of this evaluation is an average of 100 experiments: 10 experiments for each of the 10 folds starting with different random initializations. Random initialization concerns entity embeddings, weights and biases, but not word embeddings not randomly initialized, but pre-trained.

| **Entity matching:** (exact or partial) | exact | | | exact | | | partial | | | partial | | |
|---|---|---|---|---|---|---|---|---|---|---|---|---|
| **Considering hierarchy:** (yes or no) | no | | | yes | | | no | | | yes | | |
| **Metric:** | P | R | F1 ($SD_{F1}$) | P | R | F1 ($SD_{F1}$) | P | R | F1 ($SD_{F1}$) | P | R | F1 ($SD_{F1}$) |
| Chemical | 77.42 | 75.08 | 76.07 (8.52) | 77.42 | 75.08 | 76.07 (8.52) | 84.46 | 81.53 | 82.67 (7.24) | 84.46 | 81.53 | 82.67 (7.24) |
| Genomic_factor | 38.97 | 19.25 | 22.86 (16.99) | 71.58 | 71.66 | 71.41 (5.97) | 45.75 | 23.58 | 27.68 (19.15) | 83.37 | 83.51 | 83.19 (5.90) |
| ↳ Gene_or_protein | 86.10 | 85.56 | 85.72 (5.03) | 86.10 | 85.56 | 85.72 (5.03) | 91.00 | 90.40 | 90.58 (3.84) | 90.47 | 89.88 | 90.05 (3.89) |
| ↳ Genomic_variation | 25.62 | 2.48 | 2.67 (8.80) | 48.91 | 50.31 | 49.13 (8.96) | 27.09 | 3.52 | 3.83 (10.51) | 70.70 | 73.12 | 71.18 (9.55) |
| ↳ Limited_variation | 45.80 | 49.41 | 47.08 (9.13) | 45.75 | 49.35 | 47.02 (9.14) | 70.57 | 76.52 | 72.71 (9.53) | 69.46 | 75.34 | 71.57 (9.50) |
| ↳ Haplotype | 73.16 | 66.47 | 66.97 (19.01) | 73.16 | 66.47 | 66.97 (19.01) | 78.76 | 72.73 | 72.47 (19.34) | 78.76 | 72.73 | 72.47 (19.34) |
| Phenotype | 35.15 | 31.96 | 31.76 (9.09) | 51.23 | 50.95 | 50.80 (7.54) | 53.01 | 48.91 | 48.48 (10.03) | 70.09 | 69.95 | 69.57 (5.40) |
| ↳ Disease | 66.36 | 69.03 | 66.90 (8.53) | 66.34 | 69.01 | 66.88 (8.53) | 75.01 | 78.18 | 75.68 (7.50) | 71.85 | 75.07 | 72.59 (7.30) |
| ↳ Pharmacokinetic_phenotype | 38.06 | 30.33 | 29.30 (16.72) | 38.06 | 30.33 | 29.30 (16.72) | 45.81 | 38.17 | 36.47 (19.32) | 45.64 | 37.96 | 36.27 (19.40) |
| ↳ Pharmacodynamic_phenotype | 40.95 | 37.82 | 38.54 (9.80) | 40.91 | 37.78 | 38.50 (9.80) | 62.12 | 58.56 | 58.84 (9.80) | 61.43 | 57.88 | 58.18 (10.11) |
| Macro average | 52.76 | 46.74 | 49.15 (5.27) | 59.95 | 58.65 | 59.11 (6.05) | 63.36 | 57.21 | 59.76 (5.03) | 72.62 | 71.70 | 71.93 (5.64) |

Table S1: Detailed performances of the task of named entity recognition in terms of Precision (P), Recall (R), F1-score (F1) and F1-score standard deviation in brackets ($SD_{F1}$).

| **Considering hierarchy:** (yes or no) | no | | | yes | | |
|---|---|---|---|---|---|---|
| **Metric:** | P | R | F1 ($SD_{F1}$) | P | R | F1 ($SD_{F1}$) |
| isAssociatedWith | 38.75 | 28.91 | 30.89 (6.84) | 53.22 | 52.14 | 51.71 (4.02) |
| ↳ influences | 40.93 | 37.08 | 36.55 (7.35) | 52.14 | 44.64 | 46.45 (5.17) |
| ↳ causes | 70.47 | 34.36 | 41.91 (13.35) | 70.47 | 34.36 | 41.91 (13.35) |
| ↳ decreases | 33.72 | 33.59 | 29.47 (9.85) | 33.72 | 33.59 | 29.47 (9.85) |
| ↳ increases | 50.03 | 15.07 | 17.94 (15.20) | 50.03 | 15.07 | 17.94 (15.20) |
| ↳ treats | 30.17 | 70.10 | 39.97 (12.60) | 30.17 | 70.10 | 39.97 (12.60) |
| isEquivalentTo | 72.68 | 90.52 | 79.76 (7.69) | 72.68 | 90.52 | 79.76 (7.69) |
| Macro average | 48.11 | 44.23 | 45.67 (4.37) | 51.78 | 48.50 | 49.56 (4.51) |

Table S2: Detailed performances of the task of relation extraction in terms of Precision (P), Recall (R), F1-score (F1) and F1-score standard deviation in brackets ($SD_{F1}$). Note that for leaves, performances are unchanged when considering the hierarchy.
